# A centromere but not just a centromere: structure and evolution of a selfish chromosomal supergene in monkeyflowers

**DOI:** 10.64898/2026.02.23.707561

**Authors:** Evan Stark-Dykema, Findley R. Finseth, William Conner, Lila Fishman

**Author notes:** Equal contribution (Evan Stark-Dykema, Findley R. Finseth).

## Abstract

Meiotic drivers capitalize on vulnerabilities in eukaryotic meiosis and gametogenesis to gain greater-than-equal transmission to the next generation. In most plants and animals, asymmetric female meiosis is such an arena for selfishness; centromeric variants can bias their chromosome’s segregation to the one egg cell with potentially profound effects on individual fitness, population dynamics, and genome structure. However, the mechanisms and consequences of selfish centromere evolution remain obscure, due to both the complexity of centromeric DNA and to a paucity of model systems. Here, we build on representative chromosome-scale assemblies of yellow monkeyflowers *(M. guttatus* species complex) to compare the structure, gene content, and sequence of three functionally distinct haplotypes at Meiotic Drive Locus on Chromosome 11 (MDL11), whose driving *D* allele was previously shown to be centromere-containing, costly, and maintained as a balanced polymorphism in wild populations. Relative to both moderately-resistant *M. guttatus* nondrivers (*D^−^;* IM767) and weak (*d) M. nasutus*, *D* minimally evolved via two rearrangements (including a hemicentric inversion creating dual arrays of *M. guttatus* centromeric satellite *Cent728* separated by > 200 trapped genes), massive expansions of a novel MDL11-specific subfamily of satellite repeats (*Cent728_D_*) that increase *D* chromosome length by >50%, and the accumulation of >40 extra D-only genes (EDG) from diverse genomic sources in a hemizygous pericentromeric region. Under a strictly centromeric model of selfishness, the recent expansion of *Cent728_D_* is a primary candidate for generating centromere drive, either by directly altering the binding of centromeric histone CenH3 or by shifting kinetochore size or position as an epigenetic byproduct of chromosome size. However, elevated divergence in genic presence-absence variants (both in the EDG and elsewhere) and gene sequence across MDL11 strata points to strong candidates for both collusion with the *D* centromere and for interspecific (*D^−^* vs. *d*) variation in resistance. Overall, our findings support the sequential buildup of this model centromeric driver over time, reveal intriguing parallels with adaptive and gamete-killing supergenes, and provide a strong platform for further dissection of the molecular genetic basis of its dramatic deviations from Mendelian rules.

## INTRODUCTION

Meiotic drive, broadly defined as deviations from Mendelian genetic transmission that occur during gamete production, is ubiquitous across eukaryotes. Despite the name, most classic meiotic drive systems act through post-meiotic gamete-killing, often with dramatic effects on fertility, rather than by distorting chromosomal segregation *per se*. However, in plants and animals, true meiotic drive can occur during asymmetric female meiosis that result in one large egg (or megagametophyte) vs. the four smaller sperm (or microgametophytes) of male meiosis (Zwick et al. 1999; Villena and Sapienza 2001). The unequal fates of meiotic products in female create an arena for competition among homologous chromosomes: any variant that can cause a “strong” chromosome to segregate into more than its fair share of eggs gains a transmission advantage. This fundamental vulnerability, coupled with the paradoxical diversity of centromeric DNA satellites and proteins across plants and animals (Henikoff et al. 2001; Malik and Henikoff 2001), inspired the “selfish centromere model” (SCM) (Henikoff and Malik 2002; Malik and Henikoff 2002). The SCM posits that centromeres, the chromatin complexes that mediate faithful chromosomal segregation in mitosis and meiosis, routinely evolve as selfish elements, with meiotic drive by variants in the size, sequence and/or position of centromeric DNA satellite arrays creating intra-genomic conflict that drives rapid evolution of centromeric proteins (Talbert and Henikoff 2022). The foundational logic of the SCM has been empirically confirmed in taxa as disparate as mice (Robertsonian fusion/fissions: (Chmátal et al. 2014; Akera et al. 2017; Akera et al. 2019) maize (Ab10 neocentromere; (Dawe et al. 2018; Dawe 2022) and monkeyflowers (Fishman and Saunders 2008; Fishman and Kelly 2015; Finseth et al. 2021). However, the agents, mechanisms, and consequences of female meiotic drive in these model systems appear distinct, and much remains to be learned about how female meiotic drivers gain their excess transmission and how meiotic drive shapes chromosomal function, individual fitness, and genome evolution.

The Meiotic Drive Locus on Chromosome 11 (MDL11) in yellow monkeyflowers is a centromere-associated female meiotic drive locus that segregates in natural plant populations, making it ideal for understanding the evolutionary dynamics of selfish chromosomes. MDL11 was first mapped via strong transmission ratio distortion in F_2_ hybrids between the common yellow monkeyflower, *Mimulus guttatus* (IM62 line from the high-elevation annual Iron Mountain, OR population), crossed to allopatric but closely-related selfer, *M. nasutus* (SF line) (Fishman et al. 2001). Near-100% transmission of the driving *M. guttatus* allele (*D*) over *M. nasutus* (*d*) only occurred via female function in hermaphroditic F_1_ hybrids (and later backcross generations) and was not associated with female gamete loss, implicating true meiotic drive (Fishman and Willis 2005). Further, only a functional centromere can drive near-perfectly, by segregating to the egg pole in Meiosis I, while neocentromeres like Ab10 gain up to 75% transmission by distorting Meiosis II (Malik 2005). Consistent with centromeric drive, cytogenetics demonstrated that the *M. guttatus D* chromosome carries unusually large arrays of a ∼728bp satellite sequence (henceforth, *Cent728*) identified as the primary centromeric satellite across all chromosomes (Fishman and Saunders 2008). *D* drives, but relatively weakly (∼60:40), against nondriving homologues (*D^−^*) from the same *M. guttatus* population, and homozygous male and female fertility costs together maintain *D* at an intermediate equilibrium frequency (∼35-40%) in focal populations (Fishman and Saunders 2008; Fishman and Kelly 2015). Balanced polymorphism of a costly driver creates ideal conditions for the evolution of unlinked suppressors (Crow 1991; Veller 2022), and, as predicted by the SCM, one of the two *M. guttatus* copies of the Centromeric Histone (CenH3A) experienced strong selection on the same <1000 year time-scale as *D’s* recent selfish sweep and was also coincident with a weak modifier locus in mapping of unlinked suppressors (Finseth et al. 2021). These findings have established MDL11 as a rare case of the SCM in action, with costly drive by a strong centromere variant precipitating an arms race between the DNA and protein components of centromeres and also dramatically affecting individual fitness and population dynamics in the wild. However, while early genetic linkage maps and genome assemblies in monkeyflowers enabled the discovery and characterization of MDL11, the genomic components of this locus have, like most centromeres, remained largely inaccessible. Now, however, *de novo* genome assembly approaches can overcome the challenges presented by centromere satellites and other complex chromosomal regions to provide a window into the components of this unique chromosomal driver.

MDL11, while functioning as the centromere of *M. guttatus* Chromosome 11 in heterozygotes, has already shown hints that it is a more complex “selfish supergene” (Finseth et al. 2022). Unlike single-gene selfish elements that evade suppression over long time scales by constantly duplicating and inserting themselves in new genomic locations (Muirhead and Presgraves 2021; Carvalho et al. 2022), “supergene” drivers tend to originate in regions of low recombination and accumulate additional functional components and recombination suppression. Many classic meiotic drive systems, including Segregation Distorter (Navarro-Dominguez et al. 2022), t-haplotype in mouse (Kelemen et al. 2022; Runge et al. 2024; Swanepoel et al. 2025), and Ab10 neocentromere in maize (Dawe et al. 2018), are now recognized to be selfish supergenes. In these cases, drive requires the action of two functionally distinct but genetically linked components (e.g., a kinesin plus a satellite DNA knob vehicle for Ab10). Thus, inversions that lock together these components as alternative supergene alleles are key to both their birth and their elaboration and persistence as costly polymorphisms. Centromeric drive, which takes advantage of normal meiotic machinery to distort transmission without necessarily requiring other components or prompting linked antagonists, need not involve such structural or functional complexity. However, centromeres are inherently regions of low recombination, and chromosomal rearrangements may generate meiotic drive as a side effect of shifts in the position and/or size of centromere DNA arrays. In such cases, the local suppression of recombination generated by structural divergence may further promote the accumulation of genic enhancers, including variants that create a cellular environment favorable to the driving chromosome. Centromere drive in *Mimulus* is likely to have emerged via such a layered process; recent selective sweeps by the driving *D* chromosome span hundreds of genes as well as cytogenetically-visible dual *Cent728* arrays, and recombination appears entirely suppressed between *D* and alternative MDL11 haplotypes (Finseth et al. 2021). However, until now, the lack of contiguously assembled MDL11 haplotypes has prevented full characterization of *D*’s divergence in physical structure, gene content, or evolutionary history.

Here, we build on newly generated chromosome-scale assemblies of *M. guttatus* (IM62 *D*, IM767 *D^−^*) and *M. nasutus* (SF *d*) to deconvolute the MDL11 drive locus and characterize divergence among its functionally distinct haplotypes. We first infer the changes in chromosomal structure necessary to create *D’*s dual *Cent728* arrays and then characterize patterns of satellite DNA expansion and sequence evolution relative to homologous and non-homologous centromeric arrays. These approaches suggest that the driving *D* haplotype originated with a large hemicentric (*Cent728* array-breaking) inversion that overlapped with a pre-existing genic-only inversion, followed by recent expansion of a novel family of satellite repeats (*Cent728_D_*) found exclusively on Chromosome 11. We catalog patterns of sequence and gene-content divergence across the resultant MDL11 structural strata in two *D* -polymorphic populations, revealing particularly elevated *D/D^−^* sequence divergence in a core set of trapped genes moved by both inversions. Beyond the other structural differences (i.e. upstream of the first *Cent728* array in IM62) but within the *D*-swept haplotype domain, a region of <u>E</u>xtra *<u>D</u>*-only <u>G</u>enes (henceforth EDG) contains >40 gene duplicates exclusive to *D* lines. Diverse chromosomal sources for this novel pericentromeric gene content, along with variable degrees of retention and amplification post-transfer, suggest the potential for linked enhancers to accumulate amongst the hitchhikers. Overall, our findings support a model of the sequential buildup of this centromere-containing selfish supergene over time, suggesting an evolutionary history far deeper than the recent local sweeps in the populations in which it was discovered.

## METHODS

### Genome Assembly and Alignment

Chromosome-scale genome assemblies were generated for IM62 *M. guttatus* (*D*), IM767 *M. guttatus* (*D^−^*) and SF *M. nasutus* (*d*) inbred lines as described in detail elsewhere (Lovell et al. 2025). The two *M. guttatus* lines are derived from a single high-elevation annual *M. guttatus* population at Iron Mountain (IM; Oregon, USA), where *D* has recently swept to intermediate frequency, while the SF *M. nasutus* line is derived from an allopatric population near Sherars Falls (Oregon, USA). In short, each line was sequenced to high coverage (>48x) with PacBio Hi-Fi, and then contigs were assembled with HiFiAsm+HiC v0.16.1 (Cheng et al. 2021), RACON-polished with >40x Illumina (Vaser et al. 2017) and oriented/ordered to chromosome-scale with Omni-C using the JUICER v1.8.8 pipeline (Durand et al. 2016). Each genome was separately *de novo* annotated using transcriptomes of 6-8 tissues (Lovell et al. 2025). After completion of the IM62v3 genome (Lovell et al. 2025), visual re-inspection of the Omni-C contact map revealed a few likely mis-joins, so a few contigs were re-oriented or removed to generate IM62v4 (https://phytozome-next.jgi.doe.gov/info/Mguttatusvar_IM62_v4_1; Supplemental Table 1). Gene numbers were retained through the IM62v3 to IM62v4 re-ordering.

To infer major structural differences among our three focal MDL11 genotypes, we generated Chromosome 11 synteny plots using DEEPSPACE (https://github.com/jtlovell/DEEPSPACE; blkSize =2, window size = 5kb). For visual context, gene density (200kb windows) was calculated from the *de novo* annotations (Lovell et al. 2025) and *Cent728* density (50kb windows) was calculated using BLAST (Altschul et al. 1990) of a doubled IM62 consensus nucleotide sequence, with stringent parameters (E < 1 × 10^−10^; hit >700bp) (Fishman and Saunders 2008).

### Characterization of Cent728 satellite diversity genome-wide and within MDL11

We characterized the sequence-diversity of centromeric satellite DNA arrays across each of the three reference genomes IM62v4 (*D*), IM767v2 (*D^−^*), and SFv2 (*d*) using a step-wise approach. First, we detected satellite DNA regions and putative centromere locations within them by applying RepeatObserver, with default parameters (Elphinstone et al. 2025). RepeatObserver is highly (∼93%) effective at identifying experimentally-verified centromeres as regions of unusually low sequence diversity as measured by the Shannon diversity index (Elphinstone et al. 2025). We used the Shannon diversity plots from RepeatObserver (rolling window size of 500 kb windows) to identify broad genomic regions of putatively centromeric satellite DNA for each chromosome per genome (Supplemental Table S2). In all cases, these low-diversity regions also contained repeats with monomeric units similar in length to the previously identified *Cent728* satellite (Fishman and Saunders 2008). For computationally efficient characterization of *Cent728* variation within *D*, between *D* and homologous MDL11 *Cent728* arrays, and among centromeres genome-wide, we then extracted and concatenated the central *Cent728-*containing regions of all 14 chromosomes for each other genome, plus IM62 *D*, to create three pseudo-centromere genomes (IM62v4, IM767v2 + *D*, SFv2 + *D*; Supplemental Table S2). Finally, we applied StainedGlass (Vollger et al. 2022) with default parameters to generate sequence identity heatmaps capturing variation within each Chr11 *Cent728* array and divergence from the centromeric arrays of other chromosomes and accessions. In addition to the pseudo-centromere genome approach, we applied Stained Glass to pairwise comparisons of entire chromosomes: IM62v3 (all chromosome pairs) IM62v3 and SF (Chr 11), and IM62v3 and IM767 (Chr 11). The *Cent728_D_* sequence was identified as the monomer of extremely homogenous (>98% identity) arrays expanded on *D* Chr 11.

Given the quantitative difference in *Cent728_D_* repeat numbers between IM62 *D* and IM767 *D^−^*, and the possibility of greater within-population variation (particularly within the diverse *D^−^* “population” of chromosomes, we attempted to estimate the number of *Cent728_D_* copies in Illumina sequence of other IM lines with an alignment-based method. We concatenated 5 copies of the *Cent728_D_* consensus sequence as a target and aligned existing 100-150bp Illumina libraries (n = 118; (Troth et al. 2018) to it with minimap2 v. 2.28 in both paired-end and single-end mode. To validate the method, we used the PCR-free IM62, IM767, and SF libraries generated for polishing each reference genome (Lovell et al. 2025); this approach recapitulated the assembled differences best when using paired-end alignment and allowing 1 mismatch base (vs. 0 or 2) vs. the non-ambiguous consensus bases.

### Characterization of MDL11 variation within the polymorphic Iron Mountain and Cone Peak populations

To characterize recent selective sweeps and examine patterns of divergence across MDL11 strata, we used sequence of inbred lines derived from the Iron Mountain and neighboring (<1 km away) Cone Peak populations. The IM line set (n =82; 29 *D* and 53 *D^−^)* was pruned from the larger Illumina-sequenced set (Troth et al. 2018) to represent independent lineages, to have minimum coverage (median >4x across exons genome-wide) suitable for confident variant calling, and to exclude a few lines with retained heterozygosity across MDL11. The CP lines included 10 sequenced previously (Case et al. 2016; Finseth et al. 2022) plus six additional inbred lines generated via 5 generations of inbreeding and Illumina-sequenced (150 bp, paired end reads) following the same methods (see Supplemental Table S3). Due to loss of the previously sequenced CP inbred line (CP24) carrying a rare recombinant *D* haplotype missing only the EDG region (*D_short_* haplotype) we also generated a new targeted line by genotyping a large set of wild-collected CP plants at *D* and EDG-diagnostic markers (mK00858 and mK01229, respectively (Finseth et al. 2022) and identifying a wild D_short_ carrier (CP795). CP795 was selfed to generate a small set of segregating progeny and two MDL11-homozygous plants (CP795_6, CP795_7) were selected and sequenced to confirm their *D_short_* (i.e., *D*, but missing most of the EDG) genotype. One of these (CP795_7) was included in the haplotype painting, but not in population genomic analyses.

All sequences were processed through a shared pipeline, as follows. Reads were trimmed using trimmomatic v0.39 (Bolger et al. 2014) and aligned to the IM62v4 reference genome using BWA MEM v0.7.17. Reads were filtered (mapping quality > 29) using SAMtools v1.10 (Danecek et al. 2011) and PCR-duplicates removed with Genome Analysis Toolkit (GATK) v4.1.4.1 (McKenna et al. 2010; DePristo et al. 2011). For estimation of haplotype identity and divergence, we used HaplotypeCaller (GATK) to re-align around indels and emit gvcfs of all confident sites (variant and invariant) for each line, which were then assembled into vcfs for each population with the genotypegvcfs function in GATK. Because the *M. guttatus* species complex is extremely sequence diverse, compromising short-read alignment outside of genes, vcfs were restricted to exonic sites in the IM62v4 reference genome. Following Lovell et al. 2025, the vcfs were filtered using GATK’s VariantFiltration program to remove sites on the following criteria: Quality by depth < 2.0, StrandOddsRatio > 3.0, and MappingQuality < 40.0 for all sites, plus FisherStrand > 60, MQRankSum < −12.5, ReadPosRankSum <-12.5 or >12.5 for variants. In addition, to minimize the impact of mis-annotation (i.e. repetitive DNA annotated as genes) and mis-mapping, sites were further filtered with bcftools to QUAL ≥ 20, ≥ 4 reads per sample, total depth <1000, and < 20% of samples having missing data. To visualize the extent of the swept *D* haplotype at each population, we generated a vcf of all bi-allelic sites using vcftools v0.1.15 (–012 flag), and pruned it to 1:1 orthologous genes between IM62v3/v4 and IM767v2 using OrthoFinder (Emms and Kelly 2019) and also present (>10 snps/gene) in both CP and IM datasets (n = 508, spanning all of MDL11 and >150 genes in each collinear flanking region). For each gene and individual, we then calculated % identity (1-total differences/total sites) to the IM62v4 reference using a custom python script https://github.com/FishmanLab-UM/Drive25. For visualization, each population was pruned to a subset of lines representing unique haplotype-breakpoints; the full data matrix is in Supplemental Table S4.

To examine nucleotide divergence between *D* and *D^−^ M. guttatus* lines across chromosomal strata with distinct histories, we used π_4fold_ calculated in *pixy* v1.2.10.beta2 (Korunes and Samuk 2021). We set *pixy* windows to correspond to genes and identified exonic sites annotated as 4-fold degenerate in the IM62v4 genome using its gff3 annotation file and a custom script (https://github.com/tsackton/linked-selection). Analyses were restricted to Chr11 genes with 1:1 orthology and regional synteny (i.e. if an “ortholog” was not on Chr11, it was dropped) between the IM62v4 and IM767v2 references, and sufficient exonic coverage (sites > 50 and > 0.03 of gene length; total n = 1034 genes). For statistical comparisons of divergence, we defined strata: entirely outside MDL11 locus (pre- and post-MDL11 = outside) structural differences but within the bounds of the extended *D* haplotype at IM (hap), moved by both *Inv11a* & *Inv11b* (*Inv11a&b*), moved only by *Inv11b*, and moved only by *Inv11a*. The gene-wise averages for π_4fold_ divergence between *D* and *D^−^*line sets at IM and CP are provided in Supplemental Table S5. Mean genic *D* vs. *D-* divergence across strata were compared using the “Fit Model” platform in JMP 18 (SAS institute, Cary NC).

### Copy number variation across MDL11 and its sources in the extended hemizygous EDG region

To characterize *D* vs. *D^−^* divergence in gene content, we calculated the ratio of orthologues to total gene set (1:1 orthologues plus unique to IM62 and IM767 references) across all strata (Supplemental Table S6). We conducted additional analyses of the extra *D*-only genes (EDG), which consists of ∼40 genes previously identified as *D*-novel (Finseth et al. 2021) now in a contiguously-assembled region upstream of the first *Cent728* array. To identify the likely sources of IM62’s EDG duplications, we used BLASTP of each gene’s amino-acid sequence to all three references, generally retaining the top shared hit outside the EDG (e-value < 1e-30) as the source location. However, if there were several similar hits, we conservatively retained the donor location that minimized the number of duplication/transposition/insertion events while allowing for greater post-insertion gene loss within the EDG region (maximum gap = 20 genes). In cases where there was evidence of local tandem duplication (i.e. multiple hits within the EDG), we arbitrarily assigned the 5’ gene(s) in the set as the initial duplicate/transposition from outside the region. The full BLAST results for each EDG region annotated gene are provided in Supplemental Table S7.

## RESULTS AND DISCUSSION

### Novel chromosome structure, satellite DNA expansion, and gene content define a driving centromere

We use the recently published chromosome-scale genomes for the three known MDL11 functional genotypes (IM62 *M. guttatus* = driving *D*, IM767 = partly resistant *D^−^*, *SF M. nasutus* = weak *d*) except for updating IM62v3 (Lovell et al. 2025) to IM62v4 (https://phytozome-next.jgi.doe.gov/info/Mguttatusvar_IM62_v4_1). The central regions of IM767v2 and SFv2 Chr11 are each assembled into a single large contigs that contain both their entire central *Cent728* array and one of the telomeres.

MDL11 in IM62v3 and v4 spans three contigs: one containing the entire first *Cent728* array flanked by genes starting in the syntenic region prior to MDL11 and extending past the first *Cent728* array (genes Migut.11G082100-108100), a primarily genic segment (Migut.11G108200-114100), and the terminal contig flanked by the second *Cent728* array and a telomere (Migut.11G114300-139700). The v4 update notably includes re-orientation of the central gene-dense contig within MDL11 (positions 25632394 – 28070750, Table S1; this change places the few Cent728s on one end of the contig adjacent to the second large *Cent728* array on the 5’ end of the final telomere-containing contig. None of the IM62v4 Chr11 contig breakpoints corresponds to an inversion breakpoint inferred below.

There are three major structural differences between the driving IM62 *D* and functionally distinct Chromosome 11s: inversions, dramatic expansion of *Cent728* content, and novel gene content. Although the existence of each of these features was previously inferable from cytogenetics, linkage patterns, or Illumina sequence coverage, we can now fully de-convolute them. The most notable feature of *D* is its large and dual *Cent728* arrays, which total >12 Mb (> 36% of the chromosome) and, with the intervening genic region, span > 20Mb of the IM62 Chromosome 11 (Fig. 1a). Synteny analysis of genes across Chr11 points to two inversions (Fig. 1b), one of them hemicentric (*Cent728* array-breaking), as necessary to generate the IM62 *D*-specific chromosomal structure from the shared ancestral orientation found in the other accessions (Fig. 1b). Finally, just upstream of the initial *Cent728* array (i.e. outside any inversion), IM62 contains a nearly 2Mb segment (EDG) annotated as containing 40 *D*-novel genes that are paralogous to genes found elsewhere in all three genotypes, along with five interspersed genes syntenic with IM767 and SF *M. nasutus*. Together, these structural differences lock together >2/3s of IM62 Chr11 as a single driving *D* “allele” in heterozygotes with the structurally ancestral haplotypes. Below, we characterize each of these features, as well as population genomic patterns across the region, to reconstruct the evolutionary history of this unique structural variant and identify potential functional contributors to *Mimulus* centromere drive

**Figure 1.**
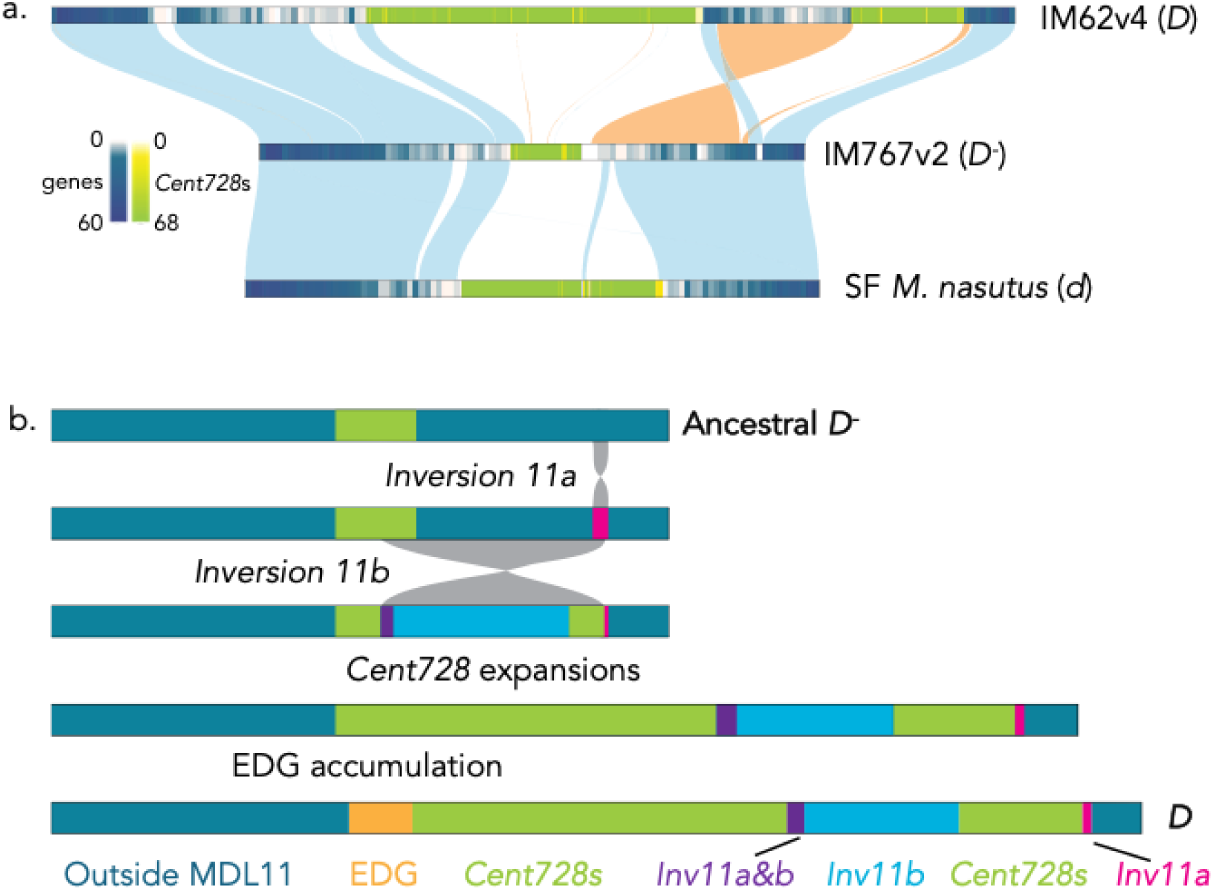
The driving IM62 *M. guttatus* Chromosome 11 is structurally distinct from both nondriving *D^−^ M. guttatus* (IM767, resistant) and weak *M. nasutus* (*d*) homologues. **a.** DeepSpace alignment of the three Chromosome 11 assemblies, showing collinear vs. inverted regions, as well as the density of genes and centromere-associated *Cent728* satellite repeats. **b.** Diagram of stepwise mutational events and resultant chromosomal strata inferred to distinguish *D* Chr11 from nondriving ancestor. We infer two overlapping inversions (genic *Inv11a* and hemicentric *Inv11b*), *Cent728* array expansions, and the build-up by duplication and transposition of the hemizygous extra *D*-only gene (EDG) region.

### Two overlapping inversions generate dual *Cent728* arrays and define genic strata within MDL11

The differences in gene order and *Cent728* distribution between the IM62 MDL11 and the three ancestral genomes are (most simply) explained by two inferred rearrangements: an initial ∼1Mb inversion *(Inv11a*) that flipped ∼90 genes on the right arm of Chr11 and a second overlapping hemicentric inversion (*Inv11b*) that trapped ∼250 genes (including most moved by *Inv11a*) between now-separated *Cent728* arrays (Fig. 1). Under this scenario, *Inv11a* generated no direct changes to centromere structure and is thus unlikely to have generated meiotic drive. However, it may have been maintained as an intermediate frequency polymorphism (like other inversions in the *M. guttatus* complex; (Veltsos et al. 2024) due to selection on either direct changes to gene function at breakpoints (i.e. disruption of protein-coding sequence or expression) or through suppression of recombination between jointly adaptive alleles. Notably, both breakpoints of *Inv11a are* associated with apparent gene loss; a transmembrane protein (MgIM767.11G103000) and a DNA-J chaperone gene (MgIM767.11G112000), both of which are highly expressed in IM767, are missing in IM62v4 relative to the other references. The exact locations of the breakpoints for *Inv11b* are less obvious; we infer the second inversion broke an ancestral *Cent728* array similar to the one found on IM767 Chr11, but the high standing diversity of *M. guttatus* makes sequence alignment outside of genes unreliable. The other *Inv11b* breakpoint occurred in the center of *Inv11a* and is not obviously associated with any gene duplication or disruption. The overlapping inversions of MDL11 *D* intriguingly parallel the complex structural divergence typical of sex chromosomes and other selfish supergenes, generating multiple strata with distinct histories of recombination suppression.

In theory, centromeric drive need not involve structural divergence outside variation in the sequence and/or abundance of CenH3-binding satellites. By changing the physical arrangement of *Cent728* arrays, the hemicentric *Inv11b* may have directly altered centromere formation even prior to array expansion. Hemicentric inversions are rarely reported in plant systems, although centromere repositioning (via multiple mechanisms) is common within and among species. Immediately following a hemicentric inversion, even when the two components cannot or do not recombine, there is the potential for formation of two independently-acting kinetochores (i.e. a dicentric chromosome) and resultant chromosomal breakage and infertility (Han et al. 2006; Fu et al. 2012). Thus, such inversions may initially be highly disruptive of both mitosis and meiosis. However, if stabilized by epigenetic inactivation of one of the now-split centromeric arrays, as appears common in experimental dicentrics (Han et al. 2006; Gao et al. 2011), the novel chromosomal arrangement may persist despite its centromere re-positioning. The feasibility of this scenario is illustrated by the (stabilized) hemicentric Chr8 of the maize knobless Tama flint line, which exhibits regional suppression of recombination (no crossovers cytogenetically observed), proper synapsis and segregation, and no elevation of infertility in F_1_ hybrids with the ancestral arrangement (Lamb et al. 2007). In addition, the breakage-fusion dynamics resulting from transient dicentrism (McClintock 1939; Birchler and Han 2018) could possibly have contributed to the duplication and movement of genes from elsewhere on Chr11 into the *D*-linked EDG region or other sites of copy number and synteny divergence (see below). Once stabilized, a hemicentric inversion may have precipitated drive through direct effects of the centromere re-positioning on kinetochore behavior, subsequent array and centromere expansion, and/or generation (e.g. by direct breakage) or recruitment of linked genic variants that facilitate drive. These stabilized scenarios of centromere silencing all predict that *D*’s functional centromere (as defined by CenH3 binding) will localize to one or the other (but not both) of its dual Cent728 satellite arrays. Future work can test these ideas using chromatin immunoprecipitation approaches to determine the location and size (see below) of functional centromere(s) in D and alternative chromosomes.

### The driving D chromosome exhibits recent expansion of a novel *Cent728* satellite (Cent728_D_)

Under the selfish centromere model, shifts in the position, size, and/or sequence of centromeric satellite DNA arrays precipitate variation in centromere “strength” that distort transmission through female meiosis. Putatively centromere-specifying *Cent728* satellites occupy a contiguous central region of each chromosome (Supplemental Table S2; (Lovell et al. 2025)) and total ∼1/7^th^ of each genome (IM62 = 16.2%, IM767 = 12.5%, Sf = 15.2%). *Cent728* content is strongly correlated with chromosome length (*r* = 0.67, *P* < 0.0001), and, with a few exceptions, *Cent728* array size and chromosome length are generally conserved across all three lines (Figure S1, Supplemental Table S2). In taxa with satellite-based centromeres, CenH3 generally localizes to tracts of highly homogeneous (i.e. recently expanded) repeats where sequence diversity is at a chromosome-wide minimum even relative to the rest of repetitive pericentromeric region generally (Elphinstone et al. 2025). In all three *Mimulus* accessions, major chromosomal minima of the Shannon diversity index always fall within the central *Cent728* arrays (Supplemental Table S2, Supplementary Figures S2-S4). However, a few other chromosomes have additional low diversity regions elsewhere. Most notably, there are highly homogeneous arrays of a shared ∼489bp satellite on Chr3 (5.16-5.71Mb on Im62v4) and Chr7 (16.23-16.7Mbin Im62v4) in all three accessions. While centromeres are defined epigenetically as the location of CenH3 binding, these consistent genome-wide patterns strongly suggest that the *M. guttatus* and *M. tilingii* species complexes have metacentric chromosomes with tandemly-repetitive regional centromeres and that *Cent728* is the predominant centromere-localized satellite.

Chr11 is a notable intra-population exception to the general conservation of chromosome lengths and correlated *Cent728* array sizes across the three *M. guttatus* complex accessions (Supplemental Fig. S1). IM62’s driving Chr11 has dramatically expanded *Cent728* content (12.18Mb, in two distinct arrays) relative to both IM767 and SF *M. nasutus*; this satellite expansion largely explains the 175% greater chromosome length of IM62 Chr11 (33.60Mb) relative to IM767 (19.06 Mb). Both RepeatObserver and StainedGlass visualizations of intra-array variation reveal highly homogeneous tracts of a specific *Cent728* subrepeat within each of *D*’s large arrays (Fig. 2, Supplemental Fig. S5-S7). This specific *D*-expanded repeat (henceforth *Cent728_D_*) occurs in large blocks with high (>98%) self-identity on the 3’ end of both *Cent728* arrays, as well as exhibiting higher order structure likely due to serial duplication of large chromosomal segments (Fig. 2b). *Cent728_D_ also* occurs in a narrow (<0.5 Mb) band in the middle of the IM767 Chr11 *Cent728* array, but is completely absent from SF *M. nasutus* and all other chromosomes in all three genomes (Fig. 2b; Supplemental Fig. S3). This pattern suggests that *Cent728_D_*was present in small amounts on an ancestral IM767-like Chr11 and then expanded dramatically subsequent to array breakage and movement by the hemicentric *Inv11b*. However, it is also theoretically possible that this sequence family originated on *D* and was subsequently adopted via gene conversion by homologous *D^−^* chromosomes. Although recombination is generally limited within centromeric arrays (and empirically nonexistent between *D* and *D^−^* MDL11 haplotypes), gene conversion is common in maize centromeric satellites (Shi et al. 2010) and a primary cause of genetic flux between alternative chromosomal arrangements in diverse systems, including our focal Iron Mountain *M. guttatus* population (Korunes and Noor 2019; Matschiner et al. 2022; Veltsos et al. 2024).

**Figure 2.**
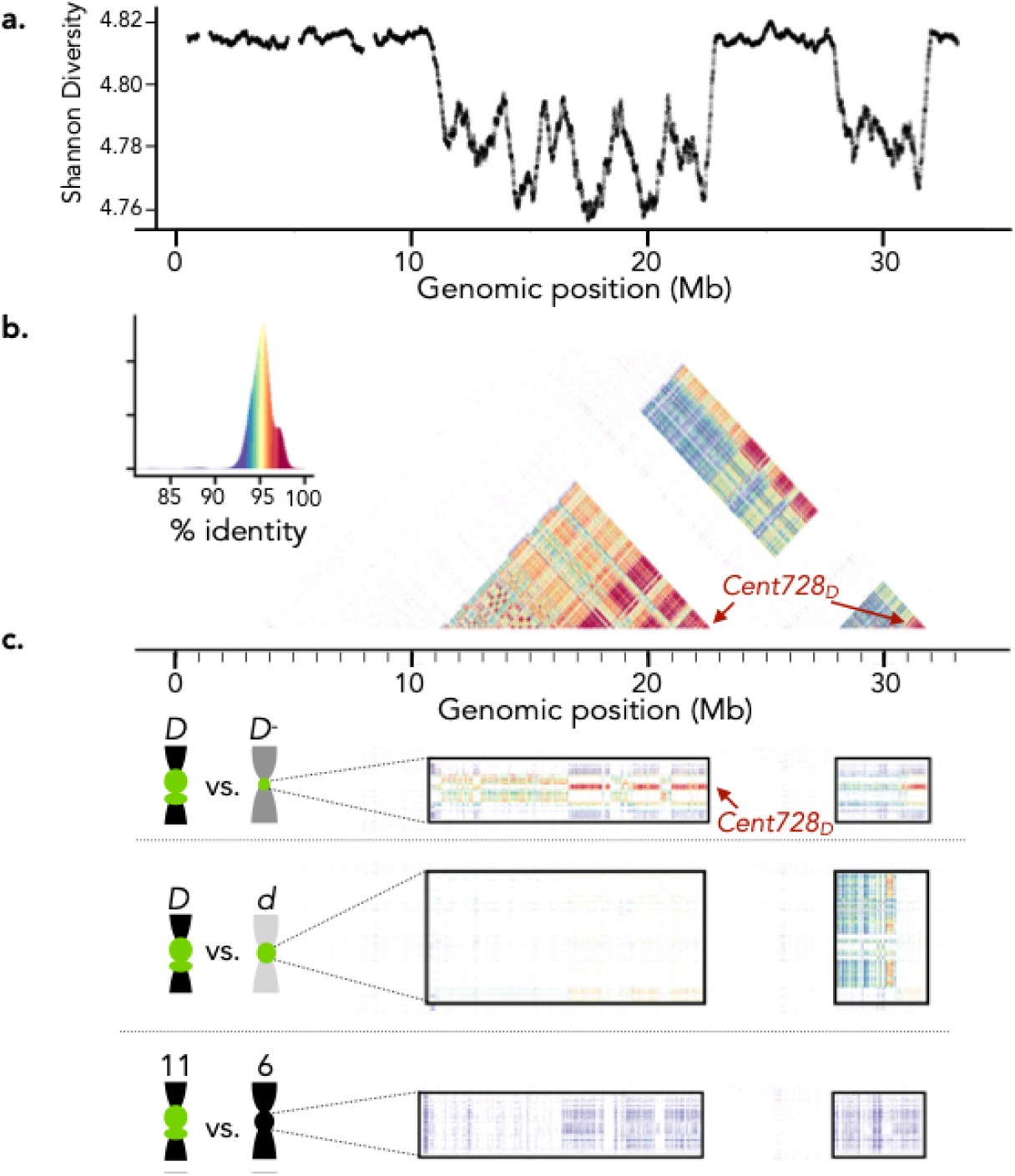
Centromere-associated *Cent728* satellites on driving IM62 *M. guttatus* Chromosome 11 (*D*) exhibit distinct sequence as well as structure and number relative to non-driving chromosomes. **a.** Shannon diversity index (from RepeatObserver) across *D* Chr11, with multiple centromere-typical diversity minima within dual *Cent728* arrays. **b.** Higher order repeat structure (intra-chromosomal identity analysis from StainedGlass) of *Cent728* arrays on *D* Chr11, showing recent expansion of homogeneous *Cent728_D_* sequences (red, arrows). **c.** Comparisons of *D* Chr11 *Cent728* satellite arrays (shared horizontal axis) to *D^−^* Chr11, *d* Chr11, and IM62 Chr6 (top to bottom on vertical axes; inter-chromosomal identities from StainedGlass), showing small region of *Cent728* (red; arrow) on *D^−^* and its absence on *d* and non-homologous chromosomes. The %identity color-scale is shared with 2b. See Supplemental Figures S5-S7 for pairwise comparisons of chromosomes among the three accessions.

Under either scenario, *Cent728_D_* expansion is a primary candidate as cause of *D*’s transmission advantage, particularly via direct effects on CenH3 binding and kinetochore formation in either homozygous or heterozygous contexts. However, while expansion of *Cent728_D_*satellites may explain drive, total *Cent728* array size does not appear to be a factor in alternative genotypes’ variable vulnerability to drive. Specifically, the relatively resistant IM767 *M. guttatus D^−^* haplotype contains far fewer total *Cent728*s (1.56 Mb, 8.2% of Chr11) than the vulnerable *M. nasut*us *d* haplotype (5.08Mb, 25.4%). This mismatch suggests that other factors, including the occurrence of *Cent728_D_* in conspecific IM767 *M. guttatus* but not SF *M. nasutus*, may contribute to stronger heterospecific drive. This association of *Cent728_D_* (but not total *Cent728*) quantity with centromere strength was not an artifact of genome (mis-)assembly, as BLAST of the PCR-free reference-polishing Illumina libraries to a multi-copy *Cent728_D_* target recapitulated its relative content in the *D*, *D^−^* and *d* reference assemblies (high, low, and none, respectively). However, this approach could unfortunately not distinguish IM62-like *D* lines (n = 44 including IM62) from *D^−^*lines (n = 76, including IM767) in more variably prepped Illumina sequences from the IM inbred line set (Troth et al. 2018), despite high homogeneity of *D* haplotypes (see below) and 76-fold amplification of *Cent728_D_* sequences in the IM62 vs. IM767 reference datasets.

A massive shift in the sequence and abundance of *Cent728_D_*, if directly accounting for drive in *M. guttatus*, would parallel the best-characterized centromere drive system, Robertsonian translocations in mice (Dudka and Lampson 2022). In mice heterozygous for fusion/fission centromeres (one metacentric vs. two acrocentric), the direction of drive reflects the relative abundance of CenH3-binding minor satellite rather than centromere arrangement *per se* (Iwata-Otsubo et al. 2017; Dudka and Lampson 2022). Strong centromeres recruit more destabilizing proteins, via distinct mechanisms in different hybrids (Akera et al. 2019), and the resultant instability in spindle attachment allows chromosomal re-orientation and thus drive. However, the occurrence of drive in mice also depends strongly on genetic background, suggesting that genic factors that modify the cellular environment of female meiosis set the stage for selfish chromosomal behavior (Kumon et al. 2021; Das et al. 2022; Walton et al. 2025). In flowering plants, female meiosis has been largely inaccessible to functional observation until very recently (Hu et al. 2025), so the molecular and cellular mechanisms underlying spindle, centromere, and kinetochore asymmetries during female meiosis remain poorly characterized. Further, despite broad conservation of function, angiosperm kinetochores only partially overlap in their protein components with the mammalian machinery – for example, of 12 proteins in the human Constitutive Centromere-Associated Network (CCAN), only one CENP-C also definitively localizes to the kinetochore in *Arabidopsis* (Pettkó-Szandtner et al. 2025). These differences, as well as the greater flexibility of the plant germ line (Böwer and Schnittger 2021) make it possible that asymmetric female meiosis in angiosperms has fundamentally different vulnerabilities to centromere drive relative to mammals. Regardless, a similarly direct mechanism for centromere strength variation in *Mimulus* would predict that the *Cent728_D_*sub-array is the specific site of centromere formation on Chr11 in Iron Mountain *M. guttatus*, and that the span of CenH3 coverage (and thus centromere size) is greater in IM62 and other *D* drivers.

Finally, while the recently expanded tracts of low diversity *Cent728_D_*within MDL11 exhibit the key hallmarks of a functional centromere location (Naish and Henderson 2024; Elphinstone et al. 2025), satellite expansions may also influence centromere strength more indirectly. In organisms with satellite centromeres, total functional centromere size (CenH3-bound chromatin) is tightly correlated with genome size (Plačková et al. 2021). Within taxa, kinetochore size variation across chromosomes is also correlated with chromosome length rather than satellite abundance and can respond plastically to changes in genomic context (Wang et al. 2021; Plačková et al. 2022). These patterns are consistent with a tension between processes that equalize CenH3 distribution across chromosomes and constraints on the minimum kinetochore size (and microtubule number) necessary to move larger chromosomes (Plačková et al. 2022). Thus, under a model in which chromosome size epigenetically determines functional centromere size, the near-doubling of Chr11 length in *D* (largely due to *Cent728* expansion) could potentially alter kinetochore formation (and thus drive) even without direct interactions between that expanded satellite sequence and centromeric histone(s). Because “fusion” products of Robertsonian translocations in mice do not consistently win meiotic contests (Chmátal et al. 2014), dramatic shifts in chromosome size cannot be a universal instigator of female meiotic drive; however, this possibility adds a distinct epigenetic mechanism for functional centromere asymmetry to the selfish centromere model. The alternative sequence-specific and sequence-independent scenarios outlined provide a clear hypothesis-testing framework for assessing the role(s) of satellite expansion on centromeric chromatin in functionally distinct MDL11 lines and their hybrids.

### MDL11 strata exhibit elevated divergence in gene content and sequence, including the hemizygous EDG region

D’s stepwise chromosomal divergence suppresses recombination and creates distinct evolutionary strata similar to those of sex chromosomes and other supergenes. This allows hitchhiking of neutral or even deleterious variants as well as the potential for the evolution of linked genic enhancers (on *D*) and resistance alleles (on *D^−^* haplotypes). D’s recent sweep to intermediate frequency in our two focal populations also generates demographic signal that extends beyond the span of structural and/or functional differences. Thus, we can define IM inbred lines (n =80) as *D* by their high genic identity to IM62 across much of Chr11 (Fig. 3). Most IM *D* haplotypes extend from ∼7.28 Mb (Migut.11G080500, ∼40 genes and 1.8 Mb before EDG structural differences) to ∼32.10Mb (Migut.11G116500, approximately co-incident with the *Inv11a* breakpoint). The two exceptions at IM (IM852 and Z485) still extend well beyond the structurally divergent EDG region to Migut.11G083200, and Migut.11G081500, respectively (Fig. 3a). Consistent with previous work, *D* haplotypes are generally shorter in the nearby Cone Peak population (Fig. 3a), where there are also rare carriers of truncated and putatively recombinant haplotypes missing most of the EDG (*D_short_*) (Finseth et al. 2022). Notably, our targeted *D_short_* line (CP795) has a distinct *D/D^−^* recombination breakpoint relative to previously sequenced *D_short_*line CP24; both breakpoints (bottom two tracks of CP *D* haplotypes; Fig. 3a) are within the small group of IM767-IM62 syntenic genes (Migut.11G087900-88200) within the EDG region, but clearly one gene apart. Distinct recombination events midway through the EDG region suggest that these rare *D_short_* haplotypes are indeed recent recombinants with *D^−^* vs. ancestral pre-EDG *D* genotypes. Unique breakpoints within the same small syntenic chunk also suggest constraints on recombination across this largely hemizygous region, along with limited spread of *D_short_* haplotypes within the subpopulation of driving chromosomes in the Cone Peak population.

**Figure 3.**
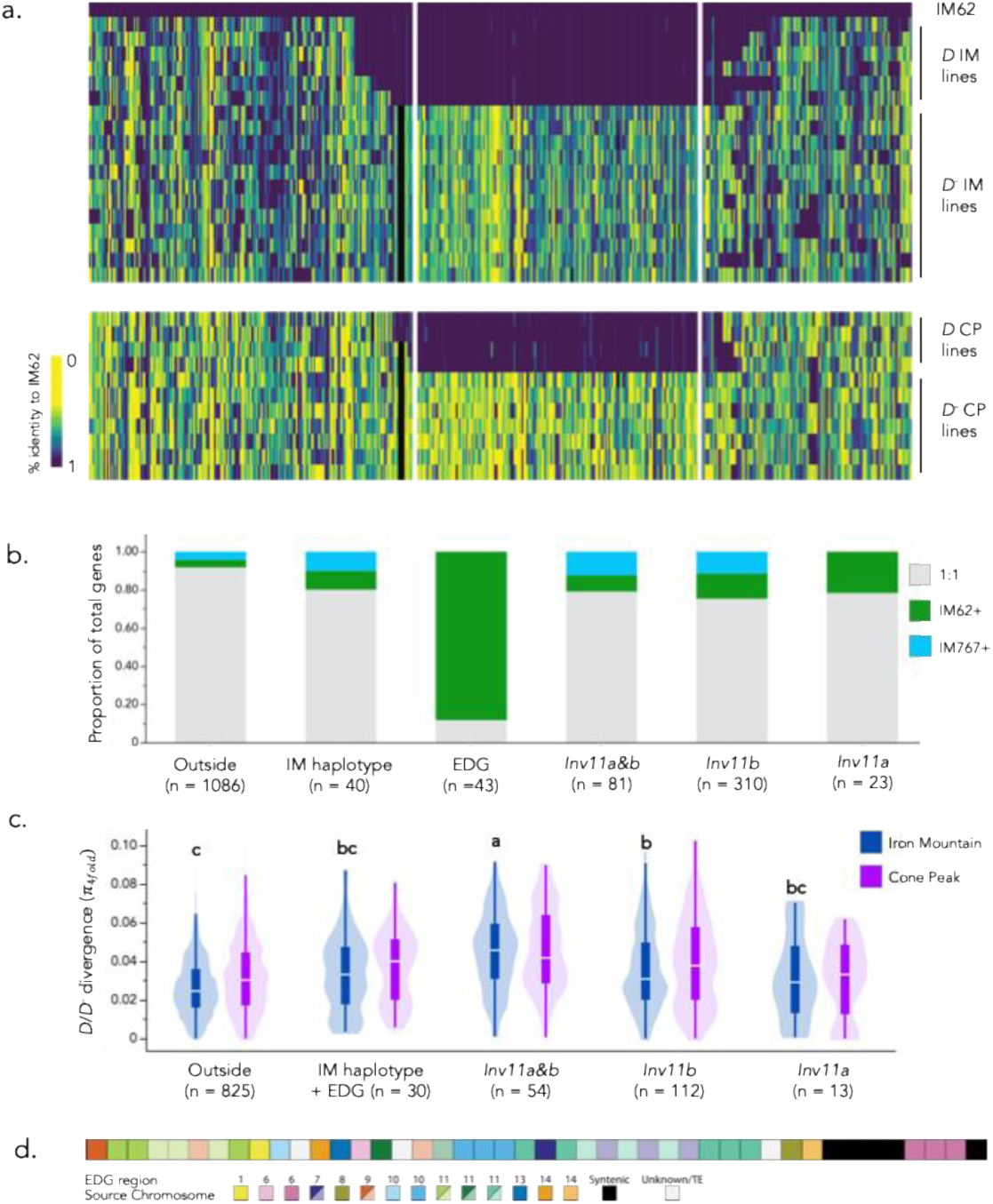
Population sweeps and elevated divergence in gene sequence and gene content across MDL11 reveals the consequences of recent selection. **a**. Representative gene-sequence haplotypes for *D* (near-identical to IM62 across MDL11) and *D*^−^ (not) inbred lines in the Iron Mountain (IM) and Cone (CP) populations of *Mimulus guttatus*. White bars indicate location of Cent728 arrays and black bar indicates the Extra D-only Genes (EDG) region. The full set of Chr11 genes on either side of the MDL11 haplotype is not shown. **b.** Gene content differences between IM62 (*D*) and IM767 (*D*^−^) reference genomes across Chr11 strata, including the largely non-syntenic EDG region just prior to the first *Cent728* array and the extended IM haplotype beyond it. Totals are the union of all genes, including orthologues (1:1) and those present in only IM62 (IM62+) or IM767 (IM767+). **c.** Average silent-site divergence (*π*_4fold_ for 1:1 orthologues only) between *D* and *D*^−^ line sets from Iron Mountain and Cone Peak populations across Chr11 strata, with the five syntenic EDG genes included in IM haplotype. Letters indicate significant differences among strata at IM from a Tukey’s HSD test. **d.** Inferred chromosomal sources of genes duplicated into the EDG region. Duplicated and transposed genes were assigned to a single source if within 10 genes of one another in putative source location. Paler shades indicate duplications (generally tandem) within the EDG.

To investigate sequence and gene content variation associated with *D’s* establishment, we examined four primarily gene-orthologous and locally syntenic strata, along with the primarily non-syntenic EDG: IM haplotype-only outside any structural differences (Fig. 3a) along with the structural distinct *Inv11a&b* and *Inv11b* between the two *Cent728* arrays, and *Inv11a*-only past the second *Cent728* array (Fig. 1b). While the EDG region (Migut.11G084400 - Migut.11G088400; ∼2Mb) shows a strongly asymmetric accumulation of genetic material (40 additional genes in IM62 *D*), and the small *Inv11a* region only has additional genes in IM62, gene content variation elsewhere in MDL11 is significantly elevated but generally symmetric (Fig. 3b). Across the non-EDG MDL11 strata (*Inv11a*, *Inv11b*, *Inv11a&b*), 21-25% of each joint gene set was not syntenic vs. only 8% (89/1086) of those in background regions entirely outside the influence of *D* (Pearson X^2^ 2 d.f. = 56.7; *P* < 0.0001) (Fig. 3b). The size and contiguity of the EDG hemizygous region, along with near-perfect identity of all *D* lines to the IM62 reference (despite previous mis-assembly), allowed us to define most of its genes as exclusive to *D* using coverage in Illumina datasets of inbred lines (Finseth et al. 2021; Finseth et al. 2022). However, even using closely related reference genomes constructed using the same assembly and annotation pipelines, inference of presence-absence variation (PAV) requires careful hand-curation of gene sets (Brůna et al. 2026). Further, the extremely high diversity of *M. guttatus* (Lovell et al. 2025) makes population-scale analyses of gene PAV and copy number variation particularly challenging. Thus, although there are several additional intriguing PAV candidates distinguishing IM62 and IM767 within the trapped genes of MDL11 (see below), we cannot yet systematically determine whether non-EDG gene content differences between IM62 and IM767 are *D/D^−^* diagnostic or idiosyncratic to this pair of lines.

Across 1:1 orthologous genes, sequence divergence between *D* and *D^−^* IM lines was strongly elevated inversions (Tukey’s HSD; P < 0.001) in those structurally bound to MDL11 by (π_4fold_ = 0.0363 ± 0.0011, n = 203) over the completely unlinked background (π_4fold_ = 0.0263 ± 0.0006, n = 829). However, finer parsing of strata across MDL11 reveals that the signal of elevated divergence in the inversion-trapped genes primarily derives from the subset of genes that were moved by both inversions to be adjacent to the larger *Cent728* array (*Inv11a&b*; mean π_4fold_= 0.0444± 0.0022, n = 54). In contrast, those moved only by *Inv11b* (0.0339 ± 0.0014, n = 125) or *Inv11a* (0.0305 ± 0.0039, n = 17) resemble the gene set in the extended (but collinear) IM *D* haplotype, which share only very recent demographic history with the rest of *D* (π_4fold_ = 0.0335 ± 0.0011, n = 30). (Fig. 3c). This suggests that the elevated divergence between *D* and *D^−^* haplotypes at IM may largely derive from the sampling effect of recent selfish sweeps rather than longer-term reproductive isolation and/or accelerated evolution of *D* variants prior to the introduction of a single haplotype our focal populations. This is consistent with previous coalescent models indicating that *D* could have arisen within the IM population, but also puts extra focus on highly divergent genes within *Inv11a&b* as candidate modifiers of drive. Greater isolation between *D* and *D^−^* IM chromosomal “populations” holds in the neighboring Cone Peak site, with mean divergence most significantly elevated in the *Inv11b* and *Inv11a&b* trapped gene sets (Fig. 3c).

In addition to abundant repetitive DNA (including expressed transposable elements) and fragmentary pseudogenized sequences, the EDG region contains 40 novel annotated genes, many of which have incomplete gene models despite substantial expression resulting in *de novo* annotation (Fig. 3d). EDG-specific paralogs derive from at least nine, and as many as thirteen, other genomic regions, though the extraordinarily high diversity of *M. guttatus* complicates source assignment with only a few reference genomes available. More than half of the EDG genes (including local tandem duplicates) appearing to have originated from Chr11 (Fig. 3d; Supplemental Table S6). These occur in two clusters; one source includes four genes derived from between Migut.11G036300 and Migut.11G038500 and the other four from Migut.11G115200 to Migut.11G116000. Within the EDG, each of these sets (assuming each was duplicated as a contiguous unit) lost more genes than were retained, gained additional single paralogs (from Chr9, Chr1, and Chr10 and from Chr7, respectively) by insertion, and experienced multiple rounds of tandem duplication to generate a complex repetitive cluster. Two cluster of three genes each appear younger, as they exactly match contiguous gene sets on other chromosomes. EDG genes Migut.11G088300-88500 are duplicates of Migut.06G190300-190500 on Chr 6; they are inserted within the five syntenic genes close to the *Cent728* array and thus not recombined off in *D_short_* haplotypes. Similarly, Migut.11G086100-86300 are duplicates of Migut.10G086500-86300. These two clearly transferred gene clusters each span ∼15kb in their source locations and the genes all retain introns and intergenic structure on Chr11. Overall, this suggests non-homologous DNA repair (likely facilitated by transposable elements in gene-flanking regions) rather than retrotransposons or pack-MULEs as the likely mechanism of gene duplication plus transposition (Krasileva 2019). As noted above, it is also possible that transient chromosomal instability precipitated by the hemicentric inversion *Inv11b* led to some of the intra-chromosomal duplications.

As a largely nonrecombining region that is often hemizygous, the EDG region resembles the Y and W chromosomes of taxa with chromosomal sex determination (Charlesworth 2016; Charlesworth 2021) as well as supernumerary B chromosomes In the long-established Y chromosomes of mammals, as well as much younger Ys in flowering plants, gene loss by deletion and the accumulation of repetitive sequences is the dominant pattern. Overall, the EDG appears similar: gaps in gene sets likely derived from contiguous sources suggest substantial degeneration of inserted gene clusters. For example, annotated genes Migut.11G084400, Migut.11G084500, and Migut.11G085000 are duplicates of Migut.11G038300, Migut.11G037200, and Migut.11G036300, respectively, suggesting the transfer of a large (>20 genes) contiguous chunk and subsequent loss of intervening sequence plus tandem duplication to generate Migut.11G084600-Migut.11G084900. This inference of a single duplication and then gene loss is supported by weakly expressed (but incomplete and not annotated) remnants corresponding to Migut.11G037000 and Migut.11G037300 in the corresponding EDG interval. This degree of gene loss parallels that of moderately degenerate plant sex chromosomes (which are generally estimated to be >0.5 million years old), suggesting a potentially much greater age for this region than the <1500 years of *D*’s recent sweeps in our focal populations (Finseth et al. 2021).

Evidence for predominant degeneration within the EDG could mean that it pre-dates *D* entirely (i.e. that this pericentromeric region existed elsewhere on the landscape prior to becoming linked with the *D*-diagnostic inversions and drive). Previous PCR-based genotyping in Oregon *M. guttatus* populations suggested that such uncoupling of EDG from *D* might be possible (Finseth et al. 2022). However, the complex structure of EDG paralogs and extreme *M. guttatus* sequence diversity undermines assignment of diagnostic alleles outside IM and CP.

In addition, like a selfish B chromosome intent only on its own transmission via the accumulation of helper genes that manipulate asymmetric mitoses (Martis et al. 2012; Chen et al. 2022; Chen et al. 2024), the non-essential EDG is universally dispensable and non-homologous, as it has evolved by accretion plus deletion rather than only degenerating from a diploid state. This means that selection against duplicated/transposed genes with deleterious effects (e.g. though increased dosage), as well as selection for retention of *D*-beneficial ones, is highly efficient and may be enough to account for both rapid accumulation and degeneration. In addition, the rarity of each *D_short_* recombinant within the *D* chromosomal population at Cone Peak suggests that the EDG may be beneficial to *D* rather than a deleterious (or even neutral) hitchhiker, as each truncated haplotype has not swept (see below). However, detailed characterization of the nucleotide and expression divergence of EDG genes from donor sequences (which may derive from beyond our few reference lines) will be necessary to reconstruct differences in their evolutionary trajectories post-duplication and transposition.

### Divergent MDL11 strata generate a large portfolio of candidates for genic collusion or conflict with *D*

While a centromere is theoretically sufficient to generate strong female meiotic drive autonomously, MDL11 also exhibits typical features of a polymorphic supergene. Supergenes are formed when inversions or other structural variants suppress recombination among alternative haplotypes (Schwander et al. 2014; Thompson and Jiggins 2014; Gutiérrez-Valencia et al. 2021; Berdan et al. 2022). Within populations, they commonly underlie multi-component alternative strategies, such as heterostyly in flowering plants (Huu et al. 2020), sexual strategies in birds (Küpper et al. 2016; Jeong et al. 2022), or mimicry patterns in butterflies (Jones et al. 2011; Joron et al. 2011). Similarly, adaptive supergenes often segregate in species occupying predictably heterogeneous environments, maintaining distinct multi-trait syndromes in the face of deleterious cross-habitat gene flow (Lowry and Willis 2010; Matschiner et al. 2022; Gompert et al. 2025; Huang et al. 2025). In either case, the key is suppression of recombination between alleles that either experience strongly divergent selection in parallel on the landscape (Rieseberg 2001; Kirkpatrick and Barton 2006) or epistatically interact such that linkage is extra-beneficial (i.e. a co-adapted gene complex: (Dobzhansky 1970).

Linkage between co-operating components has also long been recognized as a key feature of selfish meiotic drivers. Classic drive systems, such as t-haplotype in mice (Boven and Weissing 2000; Kelemen et al. 2022; Runge et al. 2024; Swanepoel et al. 2025) and Segregation Distorter (SD) in *Drosophila* (Hartl 1977; Larracuente and Presgraves 2012; Navarro-Dominguez et al. 2022) clearly evolve as selfish supergenes. The molecular mechanism of drive in these systems requires initial linkage of a poison and its antidote or of killer and nontarget alleles to prevent gametic suicide, but each has gained additional components over time. Despite no functional overlap in its drive mechanism, neocentromeric Ab10 drive also depends on linkage between satellite DNA knobs and kinesin motors, and the associated inversions foster layers of additional complexity (Dawe et al. 2018; Brady et al. 2025). These parallels accord with theoretical arguments that polymorphic drivers should favor structural variants that genetically couple drive components and successively recruit and link adjacent enhancers (Thomson and Feldman 1974; Charlesworth and Hartl 1978; Crow 1991). Thus, although *D* may not require linked genes to drive, its ∼300 genes are a substantial target for any mutations that can enhance its strength. Potential enhancers could act to directly influence centromere strength, to alter the cellular environment or timing of female meiosis to favor *D* by maximizing the “cheating window” (Walton et al. 2025), to allocate floral resources to ovules vs. pollen (as drive is strictly through female function), or to mitigate its homozygous male and female fitness costs. Conversely, variants resistant to drive (or weakening it indirectly) should be favored across (diverse and recombining) *D^−^* haplotypes, potentially accelerating *D/D^−^* divergence across MDL11.

The supernumerary genes in the EDG region are particularly strong candidates for collusion with centromeric drive, as they are otherwise dispensable. Some of the annotated genes appear broken, but our de novo annotation only includes expressed open reading frames (ORFs) and even some unannotated sequences exhibit high expression and may still be functional via RNA interference or other mechanisms). Among those annotated, several have cell division functions that makes them candidate modifiers. These include a Cyclin L (Migut.11G086100), which is part of a complex essential for meiotic synapsis in *Arabidopsis* (Zheng et al. 2014) and a histone demethylase similar to Increased Bonsai Methylation 1 (Migut.11G087700), which interacts with Chromomethylase 3 to modulate chromatin spreading from centromeres (Cheng et al. 2022; He et al. 2022). Among ampliconic genes within the EDG, a RAD60-Small Ubiquitin-like Modifier (SUMO)-like gene originating from Chr7 (Migut.07G066200) is particularly notable for its completeness and high expression in all four EDG copies (Migut.11G086500, Migut.11G086800, Migut.11G087000, Migut.11G087200). SUMOylation is an important regulator of cell cycle transitions and kinetochore stability in diverse systems (Bhagwat et al. 2021; Zhang et al. 2021; Kalidass et al. 2025). The gain of a filament-like plant protein (FPP), which is highly expressed but split into Migut.11G087400/ Migut.11G087500 due to a premature stop in the middle of a long central exon, is also intriguing. This poorly understood family of coiled coil proteins (Gindullis et al. 2002) includes TCS1, which affects microtubule stability via interactions with KCBP/ZWICHEL, a calmodulin-binding kinesin-14 (Chen et al. 2016). Notably, both the source paralog of the EDG-duplicated FPP and the *M. guttatus* homologue of ZWICHEL (Migut.11G096600), as well as a another KAC1-like kinesin-14 (Migut.11G108900) are found within the set of genes trapped by *Inv11b*. The *Inv11a&b* stratum also includes the sole *M. guttatus* homologue of the CenH3 chaperone NASP^sim^ (Goff et al. 2020; Takeuchi et al. 2024), which is more highly expressed in buds of *D* vs. *D^−^*inbred lines (Finseth et al. 2022), is in the Chr11 90% percentile for *D* vs. *D^−^* sequence divergence (π4_-fold_ = 0.050 and 0.054 at IM and CP, respectively), and contains several radical amino acid changes unique to IM62 *D* vs. other *M. guttatus* complex alleles. However, given that *D* haplotypes are near-identical and non-recombining with *D^−^,* functional manipulations will be necessary to separate any specific role(s) of captured genes in drive. Importantly for linked fitness effects, a large number of well-characterized genes for other processes are also trapped in MDL11 (e.g. homologs of the floral development gene CYCLOIDEA, the pigment producer PHYTOENE SYNTHASE, and the circadian regulator REVEILLE, among many others; Supplemental Table S5), and thus have their fates linked to *D*. Given evidence for elevated differentiation, it is thus notable that *D* is still only mildly deleterious in *DD* homozygotes, apparently does not participate in environmentally-driven fluctuating selection governing life history variation within the IM population (Kelly 2022), and exhibits no obvious epistatic associations with other inversions segregating there (Madrigal-Roca and Kelly 2025). Thus, MDL11 polymorphism in the wild provides opportunities to explore divergence during the very early stages of the accretion and spread of a selfish supergene.

The genetically separable and functionally supernumerary EDG region provides the most obvious target for additional manipulative and population genomic investigations to tease apart any direct contributions to drive or resistance from genes in each MDL11 stratum (i.e. by comparing the strength of drive and costs in *D_short_* vs. full *D* crosses). However, because resistance to drive is highly divergent between *D*^−^ *M. guttatus* lines (e.g., IM767) and SF *M. nasutus* and may also vary more subtly within IM and CP *M. guttatus* populations only recently exposed to drive, there are also opportunities to understand the mechanisms of drive from the losing side. As with the EDG, shifts in gene content provide obvious candidates for modifiers of resistance in the *D^−^* population of chromosomes. For example, the IM767 reference reveals the gain of two adjacent genes, a Walls Are Thin 1 (WAT1)-Related transporter (MgIM767.11G110400) and a T-complex protein 1 subunit delta (CCT4) Chaperonin (MgIM767.11G110500) from Chr4 relative to IM62, as well as two diverse chromosome-scale yellow monkeyflower genomes (Lovell et al. 2025). The WAT1-RELATED paralog is a fragment, but the single-exon MgIM767.11G110500 is complete and conserved (99.4% amino acid identity to source MgIM767.04G137000) and retains overall high expression. Although not well-characterized in plants, CCT chaperonins fold actin and tubulin and act as essential regulators of meiotic protein complexes in worms (Zetka et al. 2025) and mitotic checkpoint complex assembly in yeast (Kaisari et al. 2017). Because this apparently recent capture and increased dosage of a cytokinesis chaperone is unique to the IM767 reference (the only non-*D* reference from a population with active drive), it may have been retained because it modulates the strength of meiotic drive in wild *DD^−^* heterozygotes. Although additional work will be necessary to characterize such PAV across the entire *D^−^* population of chromosomes, the orthologous genes that flank the IM767 insertion in our IM62-aligned IM dataset (Migut.11G093800 and Migut.11G094100) are both in the top 2% for *D* vs. *D^−^* divergence at IM (both π_4fold_ > 0.07), potentially suggesting a local history of selection within the *D^−^* ^“^population” exposed by drive. In addition, the poorly conserved region near a *Inv11a* boundary in IM767/IM62 (between MgIM767.11G102900 and MgIM767.11G10300; Table S5) contains a highly-expressed and sequence-conserved bonus copy of the key meiotic kleisin SYNAPTIC 1 / (SYN1/ REC8) in both SF *M. nasutus* (Minas.11G101800) and outgroup LVR *M.tilingii*, while both IM *M. guttatus* lines have only single copy on Chr6 (also shared with *M. nasutus and M. tilingii).* Given the important roles of REC8 in cohesion complexes of early meiosis (Chelysheva et al. 2005), it is possible that *Inv11a-*mediated disruption of an ancestral Chr11 duplicate in the IM *M. guttatus* lineage created a window of opportunity for centromere drive, and/or that differences in SCC1 dosage contributes to the dramatic (and primarily localized to MDL11) difference in the strength of drive in *Dd* vs. *DD^−^* hybrids. Because IM767 and SF *M. nasutus* are largely collinear (Fig. 1) and MDL11 spans ∼20cM in their F_2_ hybrid (Fishman et al. 2024 Jan 27), such hypotheses about linked modifiers are amenable to testing via genetic mapping of resistance to *D* (as well as functional tests that mimic PAV effects on expression). In addition, further comparative and population genomic analyses of *non-D* accessions aligned to the IM767 and SF references can uncover the evolutionary history of these and other candidate resistance loci in populations with and without active drive.

### Conclusions

Building on new chromosome-scale genomes, we characterize the major novel components of a centromere-containing selfish supergene in yellow monkeyflowers. The driving *D* haplotype of the MDL11 locus, carried by the IM62 reference line of *Mimulus guttatus* (and >35% of Chromosome 11s in our focal populations), involves two inversions, dramatic expansions of the centromeric satellite repeat *Cent728*, accelerated divergence in the content and sequence of genes captured by the inversions, novel gene gains by duplication and transposition in the pericentromeric EDG region, and a recent selective sweep that generates linkage disequilibrium beyond the regions of structural divergence. While we cannot yet genetically separate the contributions these components to female meiotic drive and/or drive-associated fitness costs, deconvoluting this structurally complex locus provides a roadmap to candidate contributors to female meiotic drive. These findings provide novel insight into the origin of a rare exemplar of the selfish centromere model and offer intriguing parallels to adaptive and selfish supergenes in diverse system.

## Data Availability Statement

The three genome assemblies and annotations used in this work, as well as the underlying DNA and RNA sequence datasets, are publicly available through https://phytozome-next.jgi.doe.gov/. The Illumina sequence datasets of IM and CP lines are available on the NCBI Sequence Read Archive (see Table S3 for accession numbers).

## Acknowledgments

We gratefully acknowledge the contributions of J. T. Lovell, J. Jenkins, R. Walstead, , T. Bruna, K. Barry, and T. Wheeler, as well as J. H. Willis, J. K. Kelly, and A. L. Sweigart and other collaborators on the *Mimulus* JGI Community Science Project, to generating the monkeyflower genomics resources used in this study.

## Study Funding

This work was supported by NSF DEB-2344468 to LF and FRF.

## Conflict of Interest

The authors declare no conflicts of interest.

## Supplemental Table Legends

S1. Updates to the *Mimulus guttatus* IM62v3 reference genome to generate the IM62v4 reference genome.

S2. Inference of centromere array positions in reference genomes of *M. guttatus* IM62 (*D* at Meiotic Drive Locus on Chromosome 11 or MDL11), *M. guttatus* IM767 (*D^−^* at MDL11), and SF *M. nasutus* (*d* at MDL11), with Shannon Diversity minima ranges (from RepeatObserver) and *Cent728* satellite locations from (BLAST) for each chromosome. Reference sequences from the Shannon Diversity minima ranges were stitched together to create pseudo-Cent genomes: IM62, IM767 + IM62 *D*, and SF+IM62 *D* for StainedGlass analyses.

S3. Illumina sequence datasets of Iron Mountain (IM; n = 82) and Cone Peak (CP; n =17) *Mimulus guttatus* inbred lines used for population genomic analyses, including their MDL11 haplotype calls and archive information. *D/recomb* indicates lines with truncated D_short_ haplotypes (see Fig. 3a.)

S4. Identity of IM and CP inbred lines to IM62v4 reference genome (0 = all alt, 1= all ref) for IM62-IM767 orthologous Chromosome 11 genes spanning MDL11 and flanking regions (n=509). Identity was called using vcftools -- 012 flag on a biallelic SNPs vcf to convert genotypes into numerals, averaged across snps for each gene, and then inverted to indicate 100% similarity vs. difference. A subset of lines with representative breakpoints is shown in Fig. 3a.

S5. Average gene-wise divergence (π_4fold)_ between *D* and *D^−^* lines from IM and CP populations, across MDL11 strata and elsewhere on Chr11. Calculations were performed in Pixy (Korunes et al. 2021) on Chr11 genes restricted to 1:1 gene orthologs with regional synteny between IM62v3 and IM767v2 references (n=1034). Strata correspond to those in Fig 1b, plus the extended IM Haploytpe generated by recent *D* sweep.

S6. Union of all Chr11 genes in IM62v4 (n= 1394) and IM767v2 (n=1352) reference genomes, annotated (with hand-curation) as 1:1 orthologous (requiring regional synteny), additional in IM62 (IM62+), or additional in IM767 (IM767+) for identification of presence-absence and copy number variants. The “Athal” and defline columns indicate *Arabidopsis thaliana* homologues and putative gene identities/functions obtained from the published annotations.

S7. Sources of annotated genes duplicated and transposed into the pericentromeric Extra *D*-only Gene (EDG) region of IM62v4 Chr11 relative to non-*D* genomes, with information for the best peptide BLAST hit to each of the three reference annotations. We retained top hits outside of the EDG region. In cases where multiple hits had similar identities/lengths and/or there was conflict among the genomes about source genes, we chose the location that minimized the number of transposition events (up to a maximum gap of 20 genes) on the assumption that post-transposition deletions are relatively common. “Source” indicates the inferred ancestral copy, while “event” and “cluster” refer to the order and number of genes included in an insertion event. Insertions were ordered from left to right to determine the likely number of events and for tandem duplicates within EDG, the left-most copy was assigned as “source”.

## References

Akera T et al. 2017. Spindle asymmetry drives non-Mendelian chromosome segregation. Science. 358(6363):668–672. 10.1126/science.aan0092

Akera T, Trimm E, Lampson MA. 2019. Molecular Strategies of Meiotic Cheating by Selfish Centromeres. Cell. 178(5):1132–1144.e10. 10.1016/j.cell.2019.07.001

Altschul SF et al. 1990. Basic local alignment search tool. J Mol Biol. 215(3):403–410. 10.1016/s0022-2836(05)80360-2

Berdan EL et al. 2022. Genomic architecture of supergenes: connecting form and function. Philosophical Transactions Royal Soc B. 377(1856):20210192. 10.1098/rstb.2021.0192

Bhagwat NR et al. 2021. SUMO is a pervasive regulator of meiosis. eLife. 10:e57720. 10.7554/elife.57720

Birchler JA, Han F. 2018. Barbara McClintock’s Unsolved Chromosomal Mysteries: Parallels to Common Rearrangements and Karyotype Evolution. Plant Cell. 30(4):771–779. 10.1105/tpc.17.00989

Bolger AM, Lohse M, Usadel B. 2014. Trimmomatic: a flexible trimmer for Illumina sequence data. Bioinformatics. 30(15):2114–2120. 10.1093/bioinformatics/btu170

Boven M van, Weissing FJ. 2000. Evolution at the Mouse t Complex: Why is the t Haplotype Preserved as an Integral Unit? Evolution. 54(5):1795–1808 http://www.jstor.org/stable/2640677

Böwer F, Schnittger A. 2021. How to Switch from Mitosis to Meiosis: Regulation of Germline Entry in Plants. Annu Rev Genet. 55:427–452 http://pubmed.gov/34530640. 10.1146/annurev-genet-112618-043553

Brady MJ et al. 2025. Conflicting Kinesin-14s in a single chromosomal drive haplotype. GENETICS. 230(3):iyaf091. 10.1093/genetics/iyaf091

Brůna T, Sreedasyam A, Harder AM, Lovell JT. 2026. Evolutionary and methodological considerations when interpreting gene presence–absence variation in pangenomes. NAR Genom Bioinform. 8(1):lqag011. 10.1093/nargab/lqag011

Carvalho MD et al. 2022. The wtf meiotic driver gene family has unexpectedly persisted for over 100 million years. Elife. 11:e81149. 10.7554/elife.81149

Case AL, Finseth FR, Barr CM, Fishman L. 2016. Selfish evolution of cytonuclear hybrid incompatibility in Mimulus. Proc R Soc Lond B. 283(1838):20161493. 10.1098/rspb.2016.1493

Charlesworth B, Hartl DL. 1978. Population dynamics of the segregation distorter polymorphism of Drosophila melanogaster. Genetics. 89(1):171–192 http://pubmed.gov/17248828

Charlesworth D. 2016. Plant Sex Chromosomes. Annu Rev Plant Biol. 67(1):397–420. 10.1146/annurev-arplant-043015-111911

Charlesworth D. 2021. The timing of genetic degeneration of sex chromosomes. Philos Trans R Soc B. 376(1832):20200093. 10.1098/rstb.2020.0093

Chelysheva L et al. 2005. AtREC8 and AtSCC3 are essential to the monopolar orientation of the kinetochores during meiosis. J Cell Sci. 118(Pt 20):4621–4632. 10.1242/jcs.02583

Chen J et al. 2024. The genetic mechanism of B chromosome drive in rye illuminated by chromosome-scale assembly. Nat Commun. 15(1):9686. 10.1038/s41467-024-53799-w

Chen J, Birchler JA, Houben A. 2022. The non-Mendelian behavior of plant B chromosomes. Chromosom Res. 30(2–3):229–239. 10.1007/s10577-022-09687-4

Chen L et al. 2016. TCS1, a Microtubule-Binding Protein, Interacts with KCBP/ZWICHEL to Regulate Trichome Cell Shape in Arabidopsis thaliana. PLoS Genet. 12(10):e1006266. 10.1371/journal.pgen.1006266

Cheng H et al. 2021. Haplotype-resolved de novo assembly using phased assembly graphs with hifiasm. Nat Methods. 18(2):170–175. 10.1038/s41592-020-01056-5

Cheng J et al. 2022. H3K9 demethylases IBM1 and JMJ27 are required for male meiosis in Arabidopsis thaliana. N Phytol. 235(6):2252–2269. 10.1111/nph.18286

Chmátal L et al. 2014. Centromere strength provides the cell biological basis for meiotic drive and karyotype evolution in mice. Curr Biol. 24(19):2295–2300. 10.1016/j.cub.2014.08.017

Crow JF. 1991. Why is mendelian segregation so exact? Bioessays. 13(6):305–312. 10.1002/bies.950130609

Danecek P et al. 2011. The variant call format and VCFtools. Bioinformatics. 27(15):2156–2158. 10.1093/bioinformatics/btr330

Das A et al. 2022. Epigenetic, genetic and maternal effects enable stable centromere inheritance. Nat Cell Biol. 24(5):748–756. 10.1038/s41556-022-00897-w

Dawe R et al. 2018. A Kinesin-14 motor activates neocentromeres to promote meiotic drive in maize. Cell. 173(4):1–30. 10.1016/j.cell.2018.03.009

Dawe RK. 2022. The maize abnormal chromosome 10 meiotic drive haplotype: a review. Chromosome Res. 30(2–3):205–216. 10.1007/s10577-022-09693-6

DePristo MA et al. 2011. A framework for variation discovery and genotyping using next-generation DNA sequencing data. Nature Genet. 43(5):491–498. 10.1038/ng.806

Dobzhansky TG. 1970. Genetics of the Evolutionary Process. Columbia University Press. (Columbia University Press).

Dudka D, Lampson MA. 2022. Centromere drive: model systems and experimental progress. Chromosome Res. 30(2–3):187–203. 10.1007/s10577-022-09696-3

Durand NC et al. 2016. Juicer Provides a One-Click System for Analyzing Loop-Resolution Hi-C Experiments. Cell Syst. 3(1):95–98. 10.1016/j.cels.2016.07.002

Elphinstone C, Elphinstone R, Todesco M, Rieseberg LH. 2025. RepeatOBserver: Tandem Repeat Visualisation and Putative Centromere Detection. Mol Ecol Resour. e14084. 10.1111/1755-0998.14084

Emms DM, Kelly S. 2019. OrthoFinder: phylogenetic orthology inference for comparative genomics. Genome Biol. 20(1):238. 10.1186/s13059-019-1832-y

Finseth F, Brown K, Demaree A, Fishman L. 2022. Supergene potential of a selfish centromere. Philosophical Transactions Royal Soc B. 377(1856):20210208. 10.1098/rstb.2021.0208

Finseth FR, Nelson TC, Fishman L. 2021. Selfish chromosomal drive shapes recent centromeric histone evolution in monkeyflowers. PLoS Genetics. 17(4):e1009418. 10.1371/journal.pgen.1009418

Fishman L et al. 2024. Undoing the ‘nasty: dissecting touch-sensitive stigma movement (thigmonasty) and its loss in self-pollinating monkeyflowers. Biorxiv. [published online ahead of print]. 10.1101/2024.01.25.577247

Fishman L, Kelly AJ, Morgan E, Willis JH. 2001. A genetic map in the Mimulus guttatus species complex reveals transmission ratio distortion due to heterospecific interactions. Genetics. 159:1701–1716. 10.1093/genetics/159.4.1701

Fishman L, Kelly JK. 2015. Centromere-associated meiotic drive and female fitness variation in Mimulus. Evolution. 69(5):1208–1218. 10.1111/evo.12661

Fishman L, Saunders A. 2008. Centromere-associated female meiotic drive entails male fitness costs in monkeyflowers. Science. 322(5907):1559–1562. 10.1126/science.1161406

Fishman L, Willis JH. 2005. A novel meiotic drive locus almost completely distorts segregation in Mimulus (monkeyflower) hybrids. Genetics. 169(1):347–353. 10.1534/genetics.104.032789

Fu S, Gao Z, Birchler JA, Han F. 2012. Dicentric chromosome formation and epigenetics of centromere formation in plants. J Genet Genomics. 39(3):125–130. 10.1016/j.jgg.2012.01.006

Gao Z et al. 2011. Inactivation of a centromere during the formation of a translocation in maize. Chromosome Res. 19(6):755–761. 10.1007/s10577-011-9240-5

Gindullis F, Rose A, Patel S, Meier I. 2002. Four signature motifs define the first class of structurally related large coiled-coil proteins in plants. BMC Genom. 3(1):9. 10.1186/1471-2164-3-9

Goff SL et al. 2020. The H3 histone chaperone NASPSIM3 escorts CenH3 in Arabidopsis. Plant J. 101(1):71–86. 10.1111/tpj.14518

Gompert Z et al. 2025. Adaptation repeatedly uses complex structural genomic variation. Science. 388(6744):eadp3745. 10.1126/science.adp3745

Gutiérrez-Valencia J, Hughes PW, Berdan EL, Slotte T. 2021. The Genomic Architecture and Evolutionary Fates of Supergenes. Genome Biol Evol. 13(5):evab057. 10.1093/gbe/evab057

Han F, Lamb JC, Birchler JA. 2006. High frequency of centromere inactivation resulting in stable dicentric chromosomes of maize. Proc Nat Acad Sci USA. 103(9):3238–3243. 10.1073/pnas.0509650103

Hartl DL. 1977. Mechanism of a case of genetic coadaptation in populations of Drosophila melanogaster. Proc National Acad Sci. 74(1):324–328. 10.1073/pnas.74.1.324

He C et al. 2022. Histone demethylase IBM1-mediated meiocyte gene expression ensures meiotic chromosome synapsis and recombination. PLoS Genet. 18(2):e1010041. 10.1371/journal.pgen.1010041

Henikoff S, Ahmad K, Malik H. 2001. The centromere paradox: stable inheritance with rapidly evolving DNA. Science. 293(5532):1098–1102. 10.1126/science.1062939

Henikoff S, Malik HS. 2002. Centromeres: Selfish drivers. Nature. 417(6886):227–227. 10.1038/417227a

Hu B et al. 2025. A cytological framework of female meiosis in Arabidopsis. bioRxiv. 2025.08.22.671760. 10.1101/2025.08.22.671760

Huang K et al. 2025. Inversions contribute disproportionately to parallel genomic divergence in dune sunflowers. Nat Ecol Evol. 9(2):325–335. 10.1038/s41559-024-02593-4

Huu CN et al. 2020. Supergene evolution via stepwise duplications and neofunctionalization of a floral-organ identity gene. Proc Nat Acad Sci USA. 117(37):23148–23157. 10.1073/pnas.2006296117

Iwata-Otsubo A et al. 2017. Expanded Satellite Repeats Amplify a Discrete CENP-A Nucleosome Assembly Site on Chromosomes that Drive in Female Meiosis. Curr Biol. 27(15):2365–2373.e8. 10.1016/j.cub.2017.06.069

Jeong H et al. 2022. Dynamic molecular evolution of a supergene with suppressed recombination in white-throated sparrows. eLife. 11:e79387. 10.7554/elife.79387

Jones RT et al. 2011. Evolution of a mimicry supergene from a multilocus architecture. Proc R Soc Lond B. 279(1727):316–325. 10.1098/rspb.2011.0882

Joron M et al. 2011. Chromosomal rearrangements maintain a polymorphic supergene controlling butterfly mimicry. Nature. 477(7363):203–U102. 10.1038/nature10341

Kaisari S et al. 2017. Role of CCT chaperonin in the disassembly of mitotic checkpoint complexes. Proc Natl Acad Sci. 114(5):956–961. 10.1073/pnas.1620451114

Kalidass M et al. 2025. The C-terminal SUMOylation-dependent regulation of αKNL2 governs its centromere targeting and interaction with CENH3. Plant Commun. 101617. 10.1016/j.xplc.2025.101617

Kelemen RK, Elkrewi M, Lindholm AK, Vicoso B. 2022. Novel patterns of expression and recruitment of new genes on the t-haplotype, a mouse selfish chromosome. Proc Biol Sci. 289(1968):20211985. 10.1098/rspb.2021.1985

Kelly JK. 2022. The genomic scale of fluctuating selection in a natural plant population. Evol Lett. 6(6):506–521. 10.1002/evl3.308

Kirkpatrick M, Barton NH. 2006. Chromosome inversions, local adaptation and speciation. Genetics. 173(1):419–434. 10.1534/genetics.105.047985

Korunes KL, Noor MAF. 2019. Pervasive gene conversion in chromosomal inversion heterozygotes. Mol Ecol. 28(6):1302–1315. 10.1111/mec.14921

Korunes KL, Samuk K. 2021. pixy: Unbiased estimation of nucleotide diversity and divergence in the presence of missing data. Mol Ecol Resour. 21(4):1359–1368. 10.1111/1755-0998.13326

Krasileva KV. 2019. The role of transposable elements and DNA damage repair mechanisms in gene duplications and gene fusions in plant genomes. Curr Opin Plant Biol. 48:18–25. 10.1016/j.pbi.2019.01.004

Kumon T et al. 2021. Parallel pathways for recruiting effector proteins determine centromere drive and suppression. Cell. 184(19):4904–4918.e11. 10.1016/j.cell.2021.07.037

Küpper C et al. 2016. A supergene determines highly divergent male reproductive morphs in the ruff. Nature Genet. 48(1):79–83. 10.1038/ng.3443

Lamb JC, Meyer JM, Birchler JA. 2007. A hemicentric inversion in the maize line knobless Tama flint created two sites of centromeric elements and moved the kinetochore-forming region. Chromosoma. 116(3):237–247. 10.1007/s00412-007-0096-6

Larracuente AM, Presgraves DC. 2012. The selfish Segregation Distorter gene complex of Drosophila melanogaster. Genetics. 192(1):33–53. 10.1534/genetics.112.141390

Lovell JT et al. 2025. Comparative Analyses of Four Reference Genomes Reveal Exceptional Diversity and Weak Linked Selection in the Yellow Monkeyflower (Mimulus guttatus) Complex. Mol Ecol Resour. e70012. 10.1111/1755-0998.70012

Lowry DB, Willis JH. 2010. A widespread chromosomal inversion polymorphism contributes to a major life-history transition, local adaptation, and reproductive isolation. PLoS Biol. 8(9):e1000500. 10.1371/journal.pbio.1000500

Madrigal-Roca LJ, Kelly JK. 2025. Are you with me? Co-occurrence tests from community ecology can identify positive and negative epistasis between inversions in Mimulus guttatus. PLOS One. 20(4):e0321253. 10.1371/journal.pone.0321253

Malik H. 2005. Mimulus finds centromeres in the driver’s seat. Trends Ecol Evol. 20(4):151–154. 10.1016/j.tree.2005.01.014

Malik H, Henikoff S. 2001. Adaptive evolution of Cid, a centromere-specific histone in Drosophila. Genetics. 157(3):1293–1298. 10.1093/genetics/157.3.1293

Malik H, Henikoff S. 2002. Conflict begets complexity: the evolution of centromeres. Curr Opin Genet Dev. 12(6):711–718. 10.1016/s0959-437x(02)00351-9

Martis MM et al. 2012. Selfish supernumerary chromosome reveals its origin as a mosaic of host genome and organellar sequences. Proc Nat Acad Sci USA. 109(33):13343–13346. 10.1073/pnas.1204237109

Matschiner M et al. 2022. Supergene origin and maintenance in Atlantic cod. Nat Ecol Evol. 6(4):469–481. 10.1038/s41559-022-01661-x

McClintock B. 1939. The Behavior in Successive Nuclear Divisions of a Chromosome Broken at Meiosis. Proc Nat Acad Sci USA. 25(8):405–416. 10.1073/pnas.25.8.405

McKenna A et al. 2010. The Genome Analysis Toolkit: a MapReduce framework for analyzing next-generation DNA sequencing data. Genome Res. 20(9):1297–1303. 10.1101/gr.107524.110

Muirhead CA, Presgraves DC. 2021. Satellite DNA-mediated diversification of a sex-ratio meiotic drive gene family in Drosophila. Nature Ecol Evol. [published online ahead of print]. 10.1038/s41559-021-01543-8

Naish M, Henderson IR. 2024. The structure, function, and evolution of plant centromeres. Genome Res. 34(2):161–178. 10.1101/gr.278409.123

Navarro-Dominguez B et al. 2022. Epistatic selection on a selfish Segregation Distorter supergene – drive, recombination, and genetic load. Elife. 11:e78981. 10.7554/elife.78981

Pettkó-Szandtner A, Magyar Z, Komaki S. 2025. Functional framework of the kinetochore and spindle assembly checkpoint in Arabidopsis. Plant Physiol. 199(2):kiaf461. 10.1093/plphys/kiaf461

Plačková K et al. 2022. Kinetochore size scales with chromosome size in bimodal karyotypes of Agavoideae. Ann Bot. 130(1):77–84. 10.1093/aob/mcac063

Plačková K, Bureš P, Zedek F. 2021. Centromere size scales with genome size across Eukaryotes. Sci Rep-uk. 11(1):19811. 10.1038/s41598-021-99386-7

Rieseberg LH. 2001. Chromosomal rearrangements and speciation. Trends Ecol Evol. 16(7):351–358. 10.1016/s0169-5347(01)02187-5

Runge J-N, Ullrich K, Lindholm AK. 2024. De novo assembly of the selfish t supergene reveals a deleterious evolutionary trajectory. bioRxiv. 2024.09.15.613113. 10.1101/2024.09.15.613113

Schwander T, Libbrecht R, Keller L. 2014. Supergenes and complex phenotypes. Curr Biol. 24(7):R288–R294. 10.1016/j.cub.2014.01.056

Shi J et al. 2010. Widespread gene conversion in centromere cores. PLoS Biol. 8(3):e1000327. 10.1371/journal.pbio.1000327

Swanepoel CM et al. 2025. Acquisition of ampliconic sequences marks a selfish mouse t-haplotype. Nat Commun. [published online ahead of print]. 10.1038/s41467-025-66616-9

Takeuchi H, Nagahara S, Higashiyama T, Berger F. 2024. The Chaperone NASP Contributes to de Novo Deposition of the Centromeric Histone Variant CENH3 in Arabidopsis Early Embryogenesis. Plant Cell Physiol. 65(7):1135–1148. 10.1093/pcp/pcae030

Talbert P, Henikoff S. 2022. Centromere drive: chromatin conflict in meiosis. Curr Opin Genet Dev. 77:102005. 10.1016/j.gde.2022.102005

Thompson MJ, Jiggins CD. 2014. Supergenes and their role in evolution. Heredity. 113(1):1–8. 10.1038/hdy.2014.20

Thomson GJ, Feldman MW. 1974. Population genetics of modifiers of meiotic drive. II. Linkage modification in the segregation distortion system. Theor Popul Biol. 5(2):155–162. 10.1016/0040-5809(74)90038-0

Troth A et al. 2018. Selective trade-offs maintain alleles underpinning complex trait variation in plants. Science. 361(6401):475–478. 10.1126/science.aat5760

Vaser R, Sović I, Nagarajan N, Šikić M. 2017. Fast and accurate de novo genome assembly from long uncorrected reads. Genome Res. 27(5):737–746. 10.1101/gr.214270.116

Veller C. 2022. Mendel’s First Law: partisan interests and the parliament of genes. Heredity. 129(1):48–55. 10.1038/s41437-022-00545-x

Veltsos P, Madrigal-Roca LJ, Kelly JK. 2024. Testing the evolutionary theory of inversion polymorphisms in the yellow monkeyflower (Mimulus guttatus). Nat Commun. 15(1):10397. 10.1038/s41467-024-54534-1

Villena FP-M de, Sapienza C. 2001. Nonrandom segregation during meiosis: the unfairness of females. Mamm Genome. 12:331–339. 10.1007/s003350040003

Vollger MR, Kerpedjiev P, Phillippy AM, Eichler EE. 2022. StainedGlass: interactive visualization of massive tandem repeat structures with identity heatmaps. Bioinformatics. 38(7):2049–2051. 10.1093/bioinformatics/btac018

Walton RZ, Khan SJ, Yakoubi WE, Akera T. 2025. Spindle checkpoint can secure additional cheating time for selfish expanded centromeres. Curr Biol. 35(15):3687–3696.e3. 10.1016/j.cub.2025.06.056

Wang N et al. 2021. Maize centromeric chromatin scales with changes in genome size. Genetics. 217(4):iyab020. 10.1093/genetics/iyab020

Zetka M et al. 2025. A nuclear TRiC/CCT chaperonin assembles meiotic HORMAD proteins into chromosome axes competent for crossing over. Nat Commun. 16(1):9411. 10.1038/s41467-025-64403-0

Zhang J et al. 2021. The SUMO ligase MMS21 profoundly influences maize development through its impact on genome activity and stability. PLoS Genet. 17(10):e1009830. 10.1371/journal.pgen.1009830

Zheng T et al. 2014. CDKG1 protein kinase is essential for synapsis and male meiosis at high ambient temperature in Arabidopsis thaliana. Proc Nat Acad Sci USA. 111(6):2182–2187. 10.1073/pnas.1318460111

Zwick ME, Salstrom JL, Langley CH. 1999. Genetic Variation in Rates of Nondisjunction: Association of Two Naturally Occurring Polymorphisms in the Chromokinesin nod With Increased Rates of Nondisjunction in Drosophila melanogaster. Genetics. 152(4):1605–1614. 10.1093/genetics/152.4.1605

